# DNA processing by the Kaposi’s sarcoma-associated herpesvirus alkaline exonuclease SOX contributes to viral gene expression and infectious virion production

**DOI:** 10.1101/2022.09.19.508573

**Authors:** Ella Hartenian, Aaron S. Mendez, Allison L. Didychuk, Shivani Khosla, Britt A. Glaunsinger

## Abstract

Alkaline exonucleases (AE) are present in several large DNA viruses including bacteriophage λ and herpesviruses, where they play roles in viral DNA processing during genome replication. Given the genetic conservation of AEs across viruses infecting different kingdoms of life, these enzymes likely assume central roles in the lifecycles of viruses where they have yet to be well characterized. Here, we applied a structure-guided functional analysis of the bifunctional AE in the oncogenic human gammaherpesvirus Kaposi’s sarcoma-associated herpesvirus (KSHV), called SOX. In addition to identifying a preferred DNA substrate preference for SOX, we define key residues important for DNA binding and DNA processing, and how SOX activity on DNA partially overlaps with its functionally separable cleavage of mRNA. By engineering these SOX mutants into KSHV, we reveal roles for its DNase activity in viral gene expression and infectious virion production. Our results provide mechanistic insight into gammaherpesviral AE activity as well as areas of functional conservation between this mammalian virus AE and its distant relative in phage λ.

## Introduction

The oncogenic gammaherpesvirus Kaposi’s sarcoma-associated herpesvirus (KSHV) is the etiologic agent of Kaposi’s sarcoma and several B-cell lymphoproliferative diseases and is one of the most common causes of cancer in regions of Africa with a high HIV burden(1) KSHV has a ~140 kilobase linear double stranded DNA (dsDNA) genome, which is circularized upon its entry into the nucleus of an infected cell and undergoes cycles of latency and lytic replication. During latency, the viral DNA genome is tethered to the host chromatin, where it undergoes licensed ø replication by cellular replication factors during S phase (2). Viral genome amplification and progeny virion production require a switch to lytic replication, where viral genome replication is coordinated by a suite of virally encoded proteins. DNA replication is initiated at the lytic origins on the viral genome by the assembly of replication proteins including a virally encoded DNA polymerase, helicase and primase, single stranded DNA binding protein and processivity factor (3, 4). Genomes are synthesized as head-to-tail concatemers which are then cleaved into single genome lengths during packaging into the viral capsid (5).

The exact mechanism by which herpesviral DNA replicates during the lytic cycle remains unclear (e.g., rolling-circle/sigma replication, single strand annealing, ø replication or a combination of these). However, branched structures have been observed by electron microscopy that are presumably replication intermediates that must be resolved before unit length genomes are packaged (6–11). The presence of such intermediates suggests that recombination plays a role in viral genome replication, as the canonical products of a rolling circle mechanism would not be expected to generate branched structures (12–15). In KSHV and other herpesviruses such as herpes simplex virus 1 (HSV-1) and Epstein Barr virus (EBV), resolution of these branched structures involves a virally encoded alkaline exonuclease (AE) termed SOX in KSHV (UL12 in HSV-1, BGLF5 in EBV). Herpesvirus AEs are part of a larger family of PD-(D/E)-xK deoxyribonucleases that share a similar fold and active site catalytic triad and are distantly related to the phage λ exonuclease (16, 17). The DNase activity of these enzymes appears necessary to maintain viral genome integrity during replication, as mutants affecting activity or complete loss of the enzyme result in defects in virion production and decreased stability of viral DNA (6, 9, 10, 18).

In addition to processing the viral DNA genome during replication, the gammaherpesviral homologs SOX and BGLF5 also have roles in targeting mRNA, whereby cellular gene expression is dampened by accelerated messenger RNA (mRNA) degradation, termed ‘host shutoff’ (19–23). SOX endonucleolytically cleaves a broad set of RNA polymerase II transcribed RNAs and the cleaved fragments are subsequently degraded by cellular exonucleases, resulting in their effective removal from the cell (24). Biochemical studies with recombinant SOX as well as high-throughput sequencing of RNA cleavage intermediates in cells showed that SOX RNA targeting occurs at specific sites defined by a combination of RNA sequence and structural elements(24, 25). These sites incorporate a stretch of unpaired adenosines that mediate SOX binding, which facilitates its subsequent cleavage of the RNA (25). This enhanced turnover of mRNA plays a variety of roles in the gammaherpesviral lifecycle, including promoting immune evasion, regulating levels of viral gene expression, and enabling viral trafficking and replication in B cells in mice (19, 21, 22)

SOX RNase activity and the downstream consequences of RNA decay for gene regulation are well documented, but less is known about its DNase activity. Central unknown questions include the nature of the preferred DNA substrate(s) of SOX, how the dual DNase and RNase activities are coordinated by residues within and outside of the catalytic core, and what aspects of the viral lifecycle rely on SOX catalytic activity. Here we address each of these questions through an integrated structure-function analysis of the SOX protein using *in vitro* measurements with purified components as well as functional assays in SOX-expressing cells and in the context of KSHV infection. We find that several key residues required for SOX function are conserved with the exonuclease from bacteriophage λ, supporting the idea that viral DNA processing occurs via an ancient evolutionary mechanism.

## Materials and Methods

### Reagents & Biological Resources Table

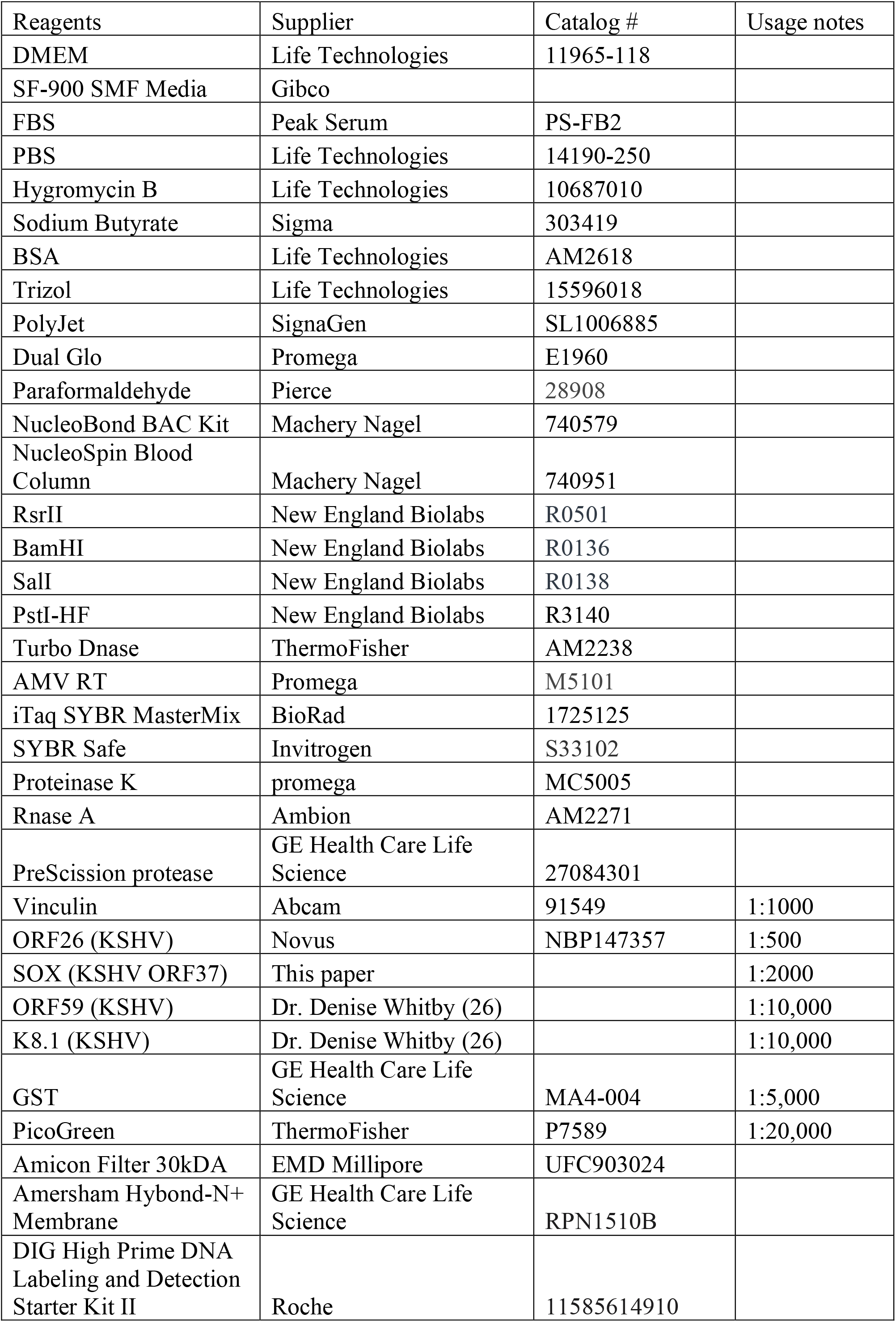
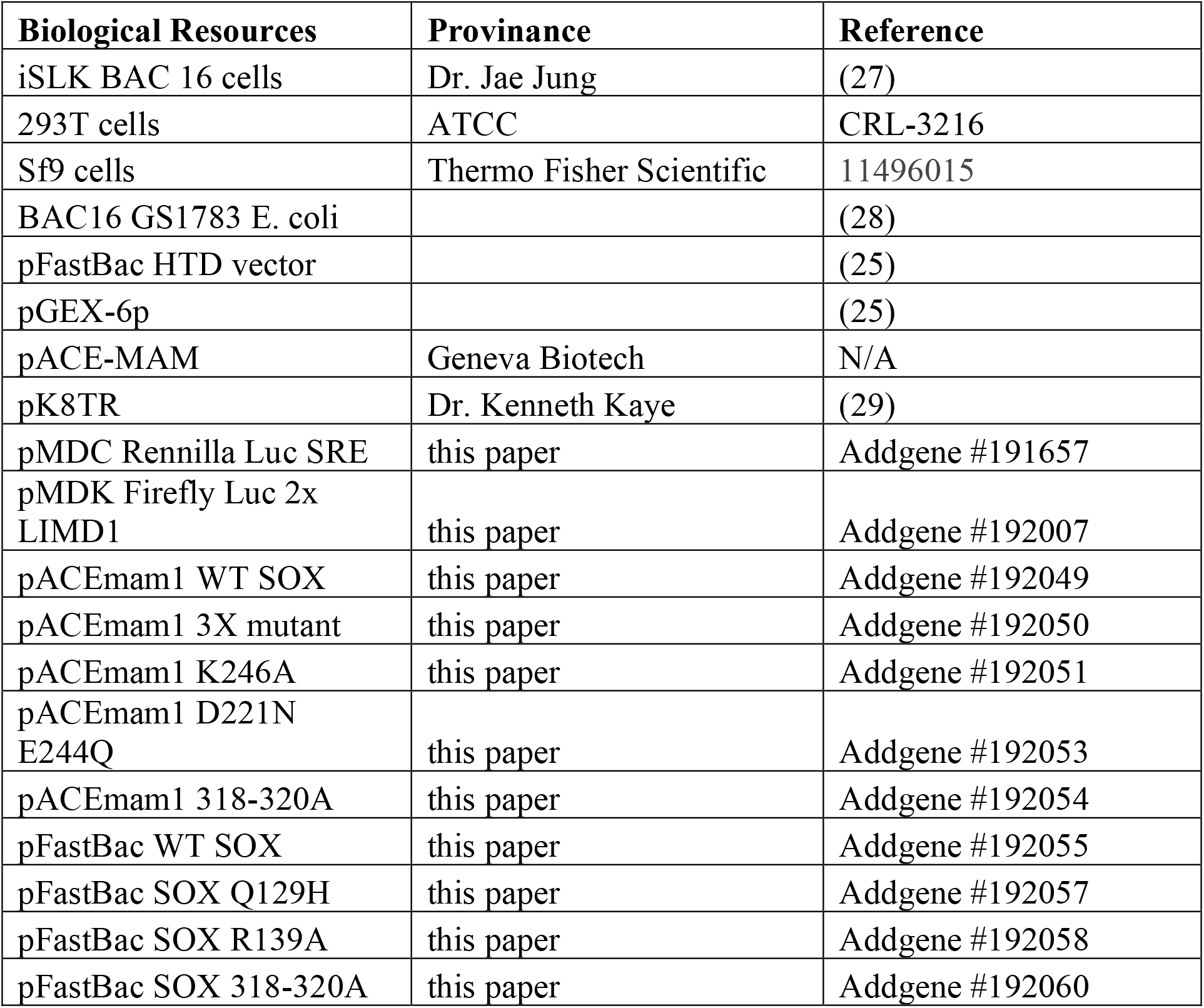

### Cloning of SOX for protein expression

KSHV SOX was codon optimized for Sf9 expression and synthesized by GENEWIZ. SOX was then subcloned into pFastBac HTD vector using restriction sites BamHI and SalI (New England Biolabs). This vector was modified to carry a GST affinity tag and PreScission protease cut site as described (25). All SOX mutants were generated using single primer site-directed mutagenesis and have been deposited in Addgene. Sequences were validated using standard pGEX forward and reverse primers. Generation of viral bacmids and transfections were prepared as described in the Bac-to-Bac® Baculovirus Expression system manual (Thermo Fisher Scientific).

### Recombinant protein expression and Purification

After transfection, Sf9 cells (Thermo Fisher Scientific) were grown for four days at 22°C using SF-900 SMF media (Gibco) substituted with 5% FBS and 1% antibiotic/antimycotic. The cellular supernatant was isolated and filtered using a 0.45 μm syringe filter and stored at 4°C away from light. In order to generate the P1 viruses, 100 μL of supernatant was transferred to a 50 mL culture of 2×10^6^ cells/mL and incubated for 96 hours. 5 mL of the P1 supernatant was transferred to a flask containing 500 mL of 2×10^6^ cells/mL and incubated for 48 hours, a time point sufficient to yield ~5 mg of SOX per 500 mL of cells. Protein expression was confirmed by western blot with and anti-GST antibody (GE Health Care Life Science).

Sf9 cell pellets were suspended in lysis buffer containing 600 mM NaCl, 5% glycerol, 0.5% Triton X-100 (Sigma), 0.5 TCEP (Denville Scientific), 20 mM HEPES pH 7.1 with a cOmplete, EDTA-Free protease inhibitor cocktail tablet (Roche). Cells were sonicated on ice using a macro tip for 3 second bursts with 17 second rests for 5 minutes at 80 amps. Cell lysates were cleared using a pre-chilled (4°C) Sorvall LYNX 6000 Superspeed centrifuge spun at 52,000xg for 30 minutes. The cleared lysate was incubated for 4 hours at 4°C using a rotating wheel with 3 mL of a GST bead slurry (GE Healthcare Life Sciences) pre-washed 3x with wash buffer (WB) containing 600 mM NaCl, 5% glycerol, 0.5 mM TCEP, 20 mM HEPES pH 7.1. The bead-protein mixture was washed 3x times with 15 mL of WB, then transferred to a 10 mL disposable column (Qiagen) and washed with an additional 50 mL of WB followed by 100 mL of low salt buffer (LSB) containing 250 mM NaCl, 5% glycerol, 0.5 mM TCEP, 20 mM HEPES pH 7.1 with periodic resuspension to prevent bead compaction. SOX was cleaved on column with PreScission protease (GE Health Life Science) overnight at 4°C, and the cleaved protein was collected with a final 8 mL LSB wash.

Cleaved protein was concentrated to ~1 mL using an Amicon Ultra-15 centrifugal filter 30 kDa MWCO (EMD Millipore), then loaded onto a HiLoad Superdex S200 pg gel filtration column (GE Healthcare Life Science). Protein elutions were concentrated using Amicon concentrators described above to ~ 10 mg/mL and 25 μL aliquots were snap frozen in liquid N_2_ using nuclease-free 0.5 mL microfuge tubes (Ambion Life Technologies) and stored at – 80°C.

### DNA substrate preparation

All DNA substrates were ordered from Integrated DNA Technologies. Sequences of DNA substrates can be found in Supplementary Table 2. All sequences contained a 3’ TAMRA fluorophore and 5’ modifications as described in Supplementary Figure 1. DNA substrates were resuspended in a distilled water to a stock concentration of 100 μM. Substrates were diluted to a working concentration of 0.1 μM using D1X buffer (50 mM NaCl and 20 mM HEPES pH 8.0). pGEX-6p (cytiva) was linearized using PstI enzyme (New England Bio labs). PstI-digested pGEX-6p was gel purified and products were cleaned up using a PCR clean up kit (Qiagen). Samples were diluted in (D1X) buffer before use in either the agarose gel or real-time DNA processing assays.

### Deoxyribonuclease assay

Observed rate constants (*k*_obs_) of SOX were determined from the cleavage kinetics of TAMRA-labeled DNA substrates as previously described (25). Briefly, 1 μL (100 nM) of DNA substrate was added to 19 μL of 25 nM SOX protein diluted in 20 mM HEPES pH 7.1, 70 mM NaCl, 2 mM MgCl_2_, 1 mM TCEP, 1% glycerol. Reactions were performed at 30°C under single turnover conditions. 3 μL of the reactions were stopped at the indicated time interval using 9 μL of STOP solution containing 95% formamide and 10 mM EDTA. Samples were resolved by 15% PAGE and imaged using a Typhoon 5 laser scanner platform (Cytiva) with imager set at Cy3 filter setting and a PMT of 625. Gels were quantified using ImageQuant and GelQuant software packages (Molecular Devices). The data were plotted and fit to exponential decay curves using Prism 9 software packages (GraphPad) to determine observed rate constants. For assays designed to detect 5’ end processing of duplex DNA, reactions were prepared as described above. Reactions were stopped by adding a buffer containing 50 μg/mL of proteinase K (Promega) in a buffer of 3 mM CaCl_2,_ 60 mM NaCl, 20 mM HEPES pH 8.0, and resolved on a 10% native page gel. Gels were imaged and quantified as described above.

### Gel-Based Exonuclease Assay

Reactions (80 uL) consisted of 250 nM PstI-linearized 5.4 kB pGEX-6p DNA (500 nM DNA ends) or 250 nM of non-linearized plasmid, 25 nM WT SOX, 20 mM HEPES (pH 7.2), 3 mM MgCl_2_, 0.5 mM TCEP, 70 mM NaCl. Each reaction was conducted at 30°C water bath. 10 μl of each reaction was removed at the indicated times and quenched by adding 2 μL 250 mM EDTA and 3 μL 6X purple loading dye (New England Bio Labs). Samples were loaded onto a 0.8% agarose gel stained with a 1/10,000 dilution of SYBR gold (Invitrogen) and visualized using a ChemiDoc MP imaging system (Bio-Rad). Control lanes labeled “ds” contain an equivalent amount of DNA as reaction lanes but not exposed to enzyme. Control lanes labeled “ss” contain half the amount of DNA heated to 95°C to mimic ssDNA products.

### Fluorescence polarization experiments (FP)

For SOX DNA binding experiments, a single-stranded DNA probe containing a 3’ TAMRA fluorophore and a 5’ phosphate. Experiments were conducted using a Spark multimode microplate reader (TECAN). The final concentration of labeled DNA was limiting (1-5 nM), and the concentration of SOX WT or mutant protein was varied in each reaction. Binding reactions of 20 μL were set up containing 20 mM HEPES-NaOH, 70 mM NaCl, 5 mM CaCl_2_, 1% glycerol, 0.2 mg/ml BSA (Ambion), 0.01% Tween 20 (Sigma-Aldrich). Reactions were allowed to reach equilibrium at room temperature for 30 minutes before the anisotropy was measured. Total fluorescence was also measured to account for quantum yield effects. The average change in anisotropy between free and bound DNA (and mean deviation) over three replicate experiments were: 0.078 ± 0.009. Multiple repeated measurements showed that equilibrium had been reached. Plotting and fitting the data to obtain *K*_D_ values was conducted assuming a “tight-binding” regime. Furthermore, saturation of binding was reached by 1 μM, so the maximum anisotropy of the complex could be directly measured. The fraction bound was then calculated and fit to the solution of a quadratic equation described an equilibrium reaction as previously reported(30). The Kd values reported are the averages and the errors of the mean deviations.

### Continuous Fluorescence-Based Exonuclease Assay

This assay was adapted from previously reports (31, 32). Reactions (25 mL) consisting of 0.01 nM PstI-linearized pGEX-6p DNA (Cytiva) (0.02 nM ends) and PicoGreen™ (1:20,000 dilution, Thermofisher) in 20 mM HEPES (pH 7.2), 3 mM MgCl_2_, 0.5 mM TCEP, 70 mM NaCl. Reactions were conducted under conditions of enzyme excess (250 nM) and fluorescence intensity was measured using a Spark multimode plate reader (TECAN) at 484 nM excitation and 522 nM emission. A reaction without enzyme served as a negative control to account for a gradual decrease in fluorescence over time, and a sample with 0.005 nM heat-denatured pGEX-6p (Cytiva) DNA (0.01 nM ssDNA) served as a positive control for ssDNA. Percent digestion were calculated from the raw fluorescence curves as described (32).

### Cells and transfections

iSLK BAC16 cells (28), kindly provided by Jae Jung’s lab, were maintained in DMEM supplemented with 10% fetal bovine serum (FBS) and 1 mg/mL hygromycin B. Uninfected iSLK-Puro cells containing a doxycycline inducible copy of the KSHV lytic transactivator ORF50 and HEK293T cells were maintained in DMEM supplemented with 10% FBS. iSLK cells were reactivated by treating with 1 μg/mL doxycycline and 1 mM sodium butyrate for 48 or 72 hours.

### Generation of the SOX Luciferase reporter

In order to generate a uniform transient transfection reporter, we utilized the MultiMam™ Transient system (Geneva Biotech). A description of our cloning strategy is as follows: SOX was cloned into an acceptor vector pACEMam1 (Geneva Biotech) using restriction sites NotI and XbaI. Firefly luciferase containing a tandem repeat of *LIMD1* within its 3’ UTR was cloned into a donor vector pMDK (Geneva Biotech) using restriction sites XhoI and KpnI. Renilla Luciferase reporter containing a 3’ UTR SOX resistance element (SRE) was cloned into doner vector pMDC (Geneva Biotech) using restriction sites XbaI and BamHI. Multigene expression constructs were generated using pACEMam1 (containing WT SOX or variants) and pMDK (Containing firefly luciferase) vectors using Cre-Lox recombination system as described in the MultiMam™ user manual. Positive clones were confirmed using full plasmid sequencing through Primordium™.

### Luciferase assay

293T cells were plated in an opaque, clear bottom 96 well plate at 1.5×10^4^ cells/well 24 hours before transfection. The cells were transfected with 10 ng of the Cre-Lox combined pMDK/pACEMam1 vector expressing luciferase and SOX and 100 ng of the pMDC vector expressing Renilla-SRE vector using PolyJet (SignaGen). The SRE is an AU-rich element that destabilizes the RNA, thus more DNA needs to be transfected to generate sufficient expression. 24 hours later media was removed, and cells were lysed in 50 μL 1x Passive Lysis Buffer (Promega) rocking at room temperature for 30 minutes. Dual Glo reagents (Promega) were used to read out firefly and renilla signal on a Tecan M1000. The firefly luminescence was normalized to the internal renilla luciferase control. The SOX-containing samples were then normalized to the empty vector control.

### RNA extraction and RT-qPCR

RNA was extracted with TRIzol followed by isopropanol purification. RNA was quantified and equal ng amounts were treated with TURBO DNase (ThermoFisher) prior to reverse transcription using AMV Reverse Transcripase (Promega) with random 9-mer primers. The resulting cDNA was quantified using iTaq Universal SYBR Mastermix (Bio-Rad laboratories) and transcript-specific primers. All qPCR data were normalized to 18S levels and the WT or vector control set to 1. PCR primer sequences are listed in Supplementary Table 2.

### Western blotting

Cells were lysed in RIPA buffer (50 mM Tris-HCl pH 7.6, 150 mM NaCl, 3 mM MgCl_2_, 10% glycerol, 0.5% NP-40, cOmplete EDTA-free Protease Inhibitors [Roche]) and then clarified by centrifugation at 21,000 x g for 10 min at 4°C. Whole cell lysate was quantified by Bradford assay and resolved by SDS-PAGE. Antibodies used for western blotting are rabbit anti-vinculin (abcam 91549, 1:1000), mouse anti-ORF26 (Novus, 1:500), rabbit anti-SOX (1:2,000), rabbit anti-ORF59 (1:10,000), and rabbit anti-K8.1 (1:10,000). The ORF59 and K8.1 antibodies are gifts from Denise Whitby (26). Purified poly clonal rabbit anti-SOX was purified in house after immunization of rabbits by YenZym antibodies, LLC.

### Generation of BAC mutants and establishing latent cell lines

The SOX Stop, Q129H, R139A, 318-20A, and D221A/E244A/K246A (3X) mutations and the corresponding mutant rescues (MR) were engineered into BAC16 using the scarless Red recombination system in GS1783 *E. coli* as described previously (33). The modified BACs were purified using the NucleoBond BAC 100 kit. BAC integrity was assessed by digestion with RsrII. The confirmed BACs were used to establish iSLK cell lines.

To generate recombinant iSLK.BAC16 cell lines, 293T cells were transfected with 5-10ug of the BAC using PolyJet (SignaGen) following the manufactures’ recommended volumes. The next day 1×10^6^ iSLK-Puro cells were plated on top of the transfected 293T cells and treated with 30 nM 12-O-Tetradecanolyphorbol-13-acdetate (TPA) and 300 nM sodium butyrate for 4 days, inducing lytic replication of KSHV. Media was then exchanged and cells were treated with media containing 300 μg/mL hygromycin B, 1 μg/mL puromycin and 250 μg/mL G418 which select for integration of the BAC and kill all the 293T cells. Over a week, the hygromycin concentration was gradually increased to a final concentration of 1 mg/mL.

### Virus characterization

To study reactivation of viral mutants, 1×10^6^ iSLK cells were plated in 10-cm dishes and induced with 1 mg/mL doxycycline and 1 mM sodium butyrate for 72 hours. To determine the fold induction of viral DNA in reactivated cells, the cells were scraped and resuspended in 10 mL PBS. 200 mL of the cell suspension was treated overnight with 80 μg/mL proteinase K (Promega) in 1x proteinase K digestion buffer (10mM Tris-HCl, pH 7.4, 100 mM NaCl, 1 mM EDTA, 0.5% SDS), after which DNA was extracted using a NucleoSpin Blood Column (Macherey Nagel). Viral DNA fold induction was quantified by qPCR using iTaq Universal SYBR Green Supermix (Bio-Rad) on a QuantStudio3 real-time PCR machine with primers for the KSHV ORF59 promoter and human CTGF promoter. Relative quantities for each sample were determined compared to a 6-point standard curve made with 1:2 dilutions of the WT reactivated sample and then ORF59 was normalized to CTGF to control for cell numbers.

Infectious virion production was determined by supernatant transfer assay. 2 mL of supernatant from a 10-cm dish of iSLK cells lytically reactivated for 72 hours (as described for gDNA harvesting) was added to 1×10^6^ 293T cells. The cell/virus suspension was spun for 2 h at 876 x *g*. After 24 hours, the medium was aspirated, the cells were washed once with phosphate-buffered saline (PBS) and cross-linked in 4% paraformaldehyde (Electron Microscopy Services) diluted in PBS for 10 minutes. The cells were washed once with PBS and analyzed on a BD Accuri 6 flow cytometer to quantify the GFP positive population.

### Viral DNA packaging assay

Southern blotting of viral genomes to monitor cleavage that occurs during viral genome packaging was performed as previously described(34). Briefly, 1×10^6^ BAC16.iSLK cells were reactivated for 72 hours, washed with PBS, and harvested by scraping. Hirt DNA extraction was performed on cell pellets followed by RNase A treatment, proteinase K treatment, phenol-chloroform extraction, and ethanol precipitation. Isolated DNA (5 μg) was digested overnight with PstI-HF (New England Biolabs) and resolved on a 0.7% 1x TBE agarose gel stained with SYBR Safe (Invitrogen) and imaged on a ChemiDoc MP to monitor total DNA. Southern blotting of the gel was performed in 20x SSC (3 M NaCl, 0.3 M sodium citrate pH 7.0) and transferred to an Amersham Hybond-N+ membrane (GE Healthcare Life Sciences) by capillary action overnight. The membrane was crosslinked in a StrataLinker 2400 (Stratagene) and DIG-labeled probe (a KSHV terminal repeat subunit derived from AscI-digested pK8TR) was hybridized, washed, and visualized with anti-DIG-AP antibody according to the DIG High Prime DNA Labeling and Detection Starter Kit II (Roche) instructions on a Chemidoc MP.

### Statistical Analyses

All statistical analysis were performed in GraphPad Prism v9.4.1 using the tests indicated in the figure legends. When comparing two samples, unpaired t-test was used, unless the comparison was to an “empty vector” control, in which case one-sample t-test was performed to account for the unequal standard deviation. When more than two samples were compared, one-way ANOVA followed by Dunnett’s multiple comparison test was used.

## Results

### SOX preferentially processes DNA templates that contain a 5’ phosphate

Studies with herpesviral alkaline exonucleases (AEs) suggest that DNase activity of these proteins contributes to viral genome processing during lytic replication (6, 11, 35, 36). However, biochemical characterization of the activity of the gammaherpesvirus AE proteins on DNA is lacking, including defining substrate preference and the extent and role of DNA binding. To identify preferred substrates for DNA processing, we incubated 25 base pair (bp) single-stranded DNA (ssDNA) or doubled-stranded DNA (dsDNA) templates derived from the KSHV LANA promoter, together with purified recombinantly expressed SOX protein (Figure 1A). Work on phage exonuclease, which is distantly related to the herpesviral AEs, has shown that dsDNA overhangs containing a 5’ phosphate are preferred substrates for efficient end processing (31). Similarly, SOX degraded substrates containing a phosphorylated 5’ 3nt overhang 21-fold (Pi 5’-3nt duplex) faster than phosphorylated blunt-ended substrates (Pi 5’-blunt) (1.1×10^5^ M^−1^/s^−1^ +/- 6.9×10^3^) versus 5.1×10^3^ M^−1^/s^−1^ +/- 2.2×10^2^) (Figure 1B, Supplementary figure 1A). Additionally, SOX cleaved ssDNA phosphate (Pi)-containing 5’ ends (ss 5’-Pi) 580-fold faster than ssDNA substrates with a 5’ hydroxyl group (SS 5’-OH) (1.08×10^6^ M^−1^/s^−1^ +/- 8.9×10^3^ versus 1.8×10^3^ M^−1^/s^−1^ +/- 3.4×10^2^) (Figure 1B, Supplementary Figure S1A). However, unlike the preference of λ exonuclease for dsDNA overhangs (37), SOX degraded phosphorylated ssDNA (SS 5’-Pi) 10-fold faster than phosphorylated 5’ dsDNA overhangs (Pi 5’-3nt duplex). The dsDNA substrate with a blunt ended 5’ hydroxyl group could not be processed by SOX (OH Blunt), suggesting that end processing of 5’ phosphate containing DNA substrates is the preferred mechanism of degrading DNA (Supplementary Figure S1A). We next monitored the activity of SOX by comparing its activity on linearized versus non-linearized plasmid. At a limiting concentration of SOX (25 nM) we observed a gradual reduction in linearized plasmid, whereas no degradation products were observed with the non-linearized plasmid in the time frame of this assay (Figure 1C). Interestingly, we observed collapse of the double stranded linearized plasmid to levels mimicking that of our ssDNA control (Figure 1C). This suggests that SOX can fully digest one strand of a linearized dsDNA substrate to produce a single-stranded intermediate. SOX displayed equivalent DNase activity on a 25 bp ssDNA template derived from a viral sequence or the same sequence scrambled (Supplementary Figure S1B). This agrees with data from other herpesvirus AEs showing that processing is not dependent on sequence features (6, 35, 36).

**Figure 1.**
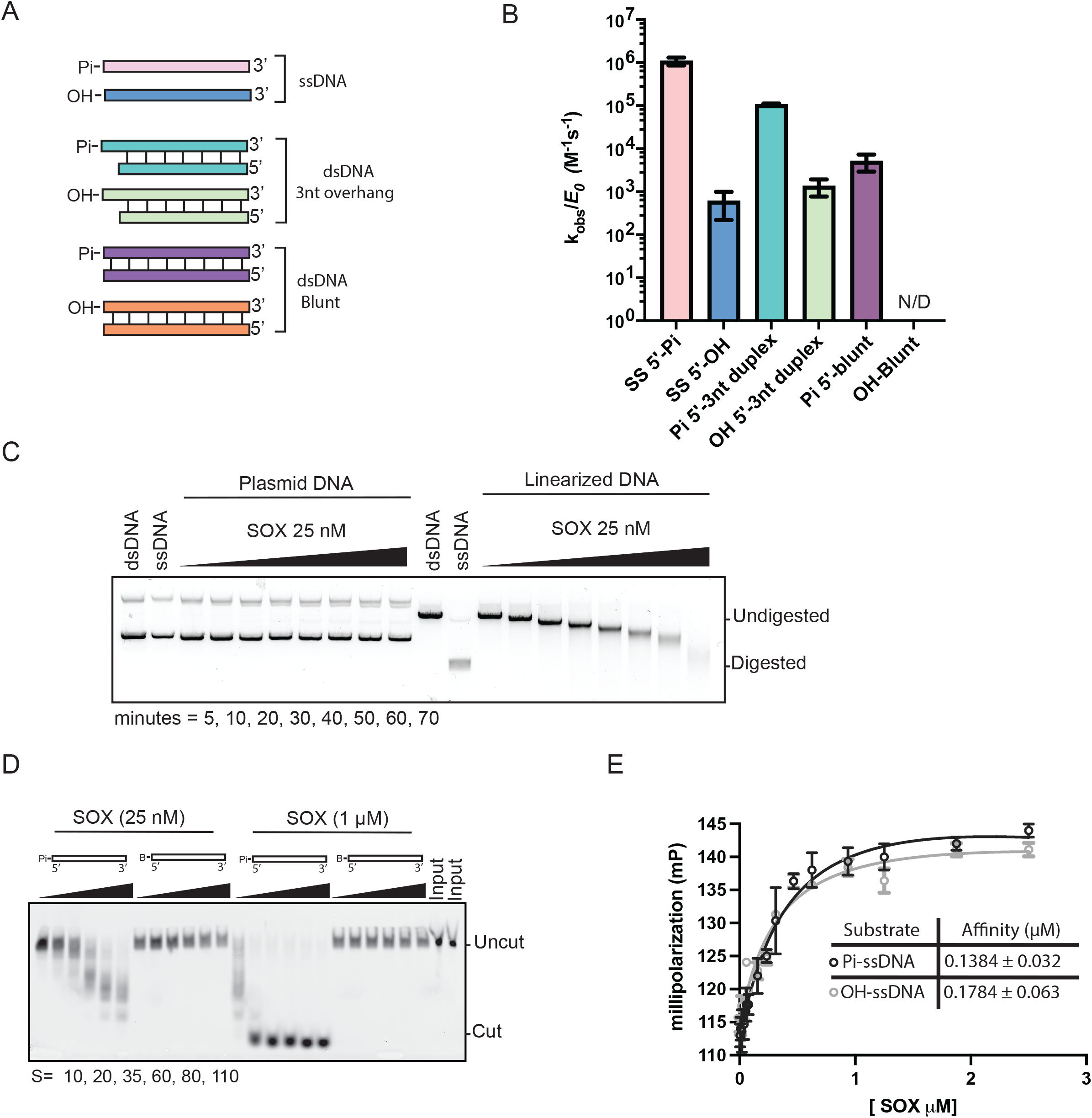
Characterization of SOX DNA exonuclease activity and DNA binding. (A) Diagram of the 25 bp LANA substrates used in DNA processing and binding experiments (Pi, 5’ phosphate; OH 5’ hydroxyl group). All substrates contain a 3’ TAMRA fluorophore. (B) Measurements of the catalytic efficiency of SOX processing of dsDNA and ssDNA substrates with different 5’ end modifications. All reactions were performed in a single turnover regime with 25 nM of SOX protein. Experiments were performed in triplicate. (C) Agarose gel-based exonuclease assays. 25 nM of WT enzyme was incubated with 250 nM linear or non-linearized pGEX-6p DNA at 30°C. Aliquots removed at the indicated times were analyzed by gel electrophoresis. Control lanes labeled ds and ss mark the substrate and product of the reaction. (D)Urea-PAGE gel showing SOX processing of ssDNA substrates with a 5’ phosphate or a 5’ fluorescein dye. Reactions were performed at a constant concentration of 25 nM and 1 μM of SOX enzyme, respectively. Aliquots removed at the indicated times were analyzed by urea-PAGE electrophoresis. (E) Fluorescence polarization binding curves using ssDNA with a 5’ phosphate and 5’hydroxyl group. Raw millipolarization units were plotted as a function of SOX concentration. Curves were fit to a single binding model for three independent measurements.

Two additional observations indicate that 5’ exonuclease rather than endonuclease activity is the primary mode of SOX DNA processing. First, SOX was unable to process a ssDNA substrate containing a bulky ‘blocking’ fluorescein dye conjugated to the 5’ end (5’-B-DNA), even in conditions containing a 40-fold excess of enzyme (Figure 1D). Second, no processing was observed on a ssDNA substrate with an unblocked 5’ end but containing a thiophosphate at positions 1 and 2, further suggesting that base hydrolysis occurs though a similar mechanism as PD/ExK exonucleases (Supplementary Figure S1C). Notably, fluorescence polarization (FP) assays showed no difference in SOX binding affinity to 5’ phosphorylated versus hydroxyl containing ssDNA (Figure 1E). Thus, while engagement of the 5’ phosphorylated end is necessary for DNA processing by SOX, it is dispensable for the initial substrate binding(36).

### Structure and functional analysis of SOX DNA binding and processing

In order to identify SOX residues critical for DNA binding and processing, we leveraged biochemical and structural work on phage λ exonuclease as well as a co-crystal structure of SOX bound to dsDNA (31, 38)(Supplementary Figure S2A,S2B). We used sequence alignments to identify conserved residues and generated 8 mutants targeting residues or groups of residues in SOX that we hypothesized would mostly likely be involved in its activity on DNA (Figure 2A, Supplementary Figure S2A). These include putative DNA binding mutants (K250A, R370A, 318-20A), as well as putative DNA processing mutants located in conserved regions either within (K246A, D221N/E244Q, D221A/E244A/K246A) or outside (Q129H, R139A) the catalytic triad (Supplementary Figure S2B,C).

**Figure 2.**
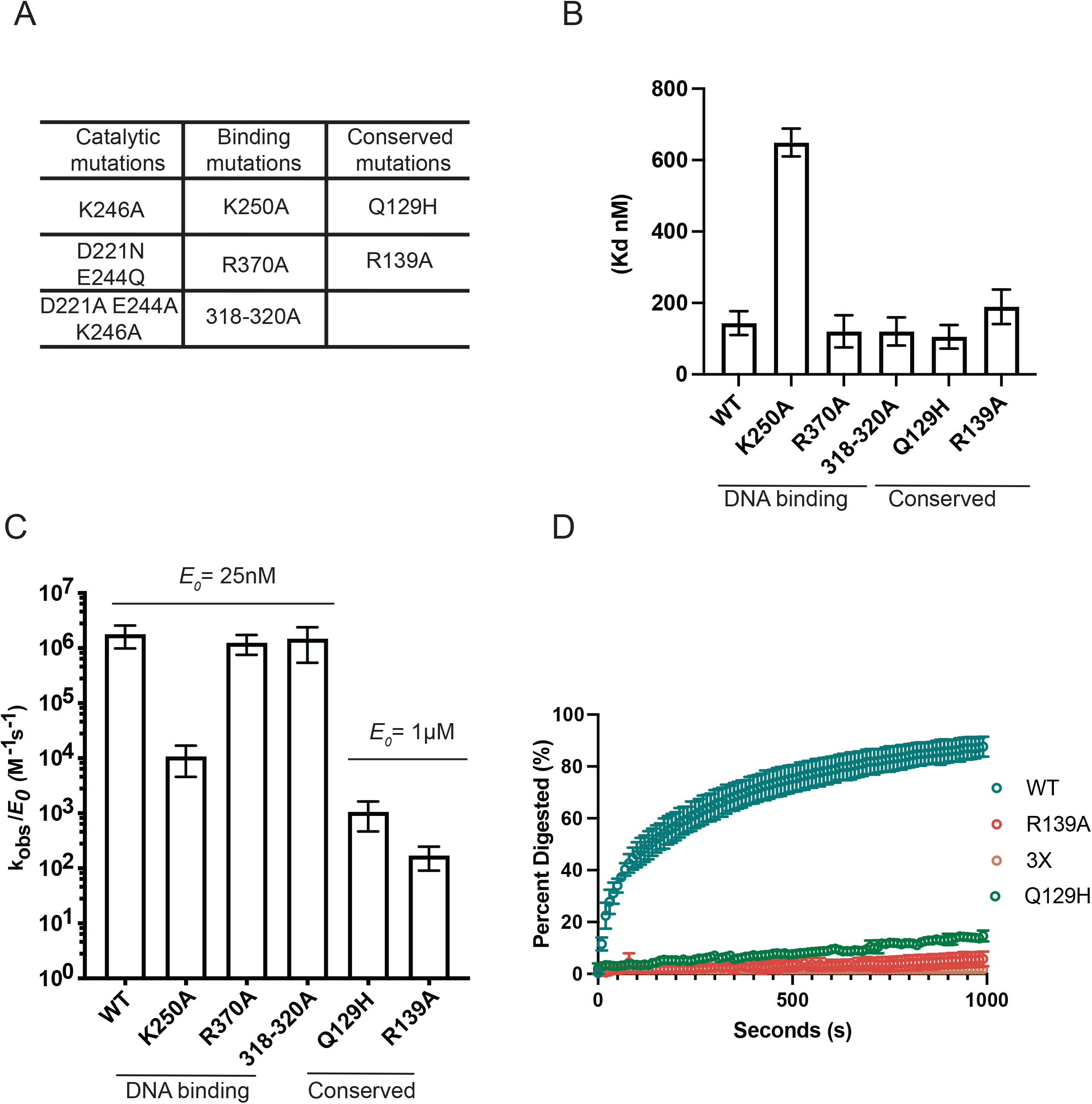
In vitro characterization of SOX DNA processing mutants. (A) A total of 8 SOX mutants were grouped into three categories: catalytic residues, putative DNA binding residues, and other highly conserved residues. (B) SOX mutant Kd values were grouped according to their putative functions and plotted. Kd and error bars were derived from fitting graphs to single binding models for three independent measurements. (C) Catalytic efficiencies were determined for all 8 SOX mutants (25 nM) in the presence of a 25 bp ssDNA probe derived from the viral LANA gene containing a 5’ phosphate and a 3’ fluorophore. Experiments were performed in triplicate. (D) The exonuclease activity of WT and mutated versions of SOX were measured using a continuous fluorescence assay. Excess amounts of enzyme was used (250 nM) in limiting amounts of substrate (0.01 nM) and plotted using percent digestion as a function of time.

We first evaluated the ability of the putative DNA binding mutants and the conserved mutants located outside of the catalytic core to bind DNA using a FP binding assay. Among these, only the K250A resulted in a substantial reduction in DNA binding (649.12 ± 39.25nM) (Figure 2B and Supplementary Figure S4). Surprisingly, although residues 318-320 and R370 appear to make contacts with the DNA backbone in the SOX-DNA crystal structure, mutation to alanine does not impair binding in our assay. These contacts could be a result of a non-productive binding interaction of SOX with DNA(36).

We next measured the cleavage kinetics of these mutants on a 5’ phosphorylated ssDNA substrate. As expected, the single and double catalytic mutants (D221N/E244Q, D221S) and triple mutant D221A/E244A/K246A (3X mutant), along with the catalytic lysine (K246A) mutant were unable to process the ssDNA substrates, validating their role in catalysis (Supplementary Figures S3A, S3B).

Outside of the catalytic triad, we also observed a loss in DNase activity with the Q129H and R139A mutants (which retain WT levels of DNA binding) using 25 nM of enzyme (Figure 2C, Supplementary Figure S3B). However, with a 40-fold excess of enzyme and a longer time course, a low level of residual activity as well as processing intermediates were detectable, suggesting that residues R139 and Q129 play a role in SOX processivity (Supplementary Figure S3C, and Supplementary Table 1). Indeed, a similar processing defect was observed with mutations at residue R28 in lambda exonuclease mutant, which is analogous to SOX residue R139 (31). Mutants R370A and 318-20A displayed no defects in catalytic activity (Figure 2C and Supplementary Table 1), in agreement with their retention of DNA binding activity, while mutant K250A, which was impaired for DNA binding, displayed a 165-fold decrease in activity relative to WT (Figure 2C and Supplementary Figure S3D).

Finally, we used a fluorescence-based continuous DNA exonuclease assay to monitor end processing in real-time and directly compare enzyme processivity of the mutants (32, 36). In this assay, a linearized 5.4 kb plasmid was digested to near completion by an excess of WT SOX (Figure 2D). In agreement with the gel-based ssDNA assay, both end processing mutants R139A and Q129H showed marked defects in their ability to process DNA and the 3X catalytic mutant showed no activity (Figure 2D). Collectively, these data delineate the importance of individual SOX residues in DNA binding (K250), enzyme processivity (Q129, R139) and catalysis (D221, E244, K246) and show that each of these activities contribute to distinct steps in processing of DNA substrates.

### Residues Q129 and R139 are required for SOX activity on DNA but not mRNA

We next examined how these SOX mutants impact its mRNA degradation activity. Although SOX does not degrade plasmid DNA in cells, it does readily degrade endogenous cellular and reporter-derived mRNA (20). We therefore established a reporter system that uses the levels of luciferase protein luminescence as a readout of SOX RNase activity in cells. Unlike its activity on DNA, SOX targets RNA for endonucleolytic cleavage at specific locations that are influenced by RNA sequence and structure (25, 39). We generated a firefly luciferase construct containing one or two copies of the well-characterized SOX targeting sequence from the cellular *LIMD1* transcript in its 3’ UTR (24, 25). This rendered the mRNA a preferred substrate for SOX cleavage, resulting in reduced firefly luciferase protein levels upon co-transfection with SOX in 293T cells (Figure 3A-B). As a transfection normalization control, we also included a renilla luciferase transcript with an RNA element derived from the interleukin-6 transcript termed a ‘SOX resistance element’ (SRE) previously shown to block SOX targeting (40, 41). Given that we observed maximal reduction in firefly luciferase levels with the construct containing two copies of the SOX targeting element (Figure 3B), we used this construct for subsequent assays.

**Figure 3.**
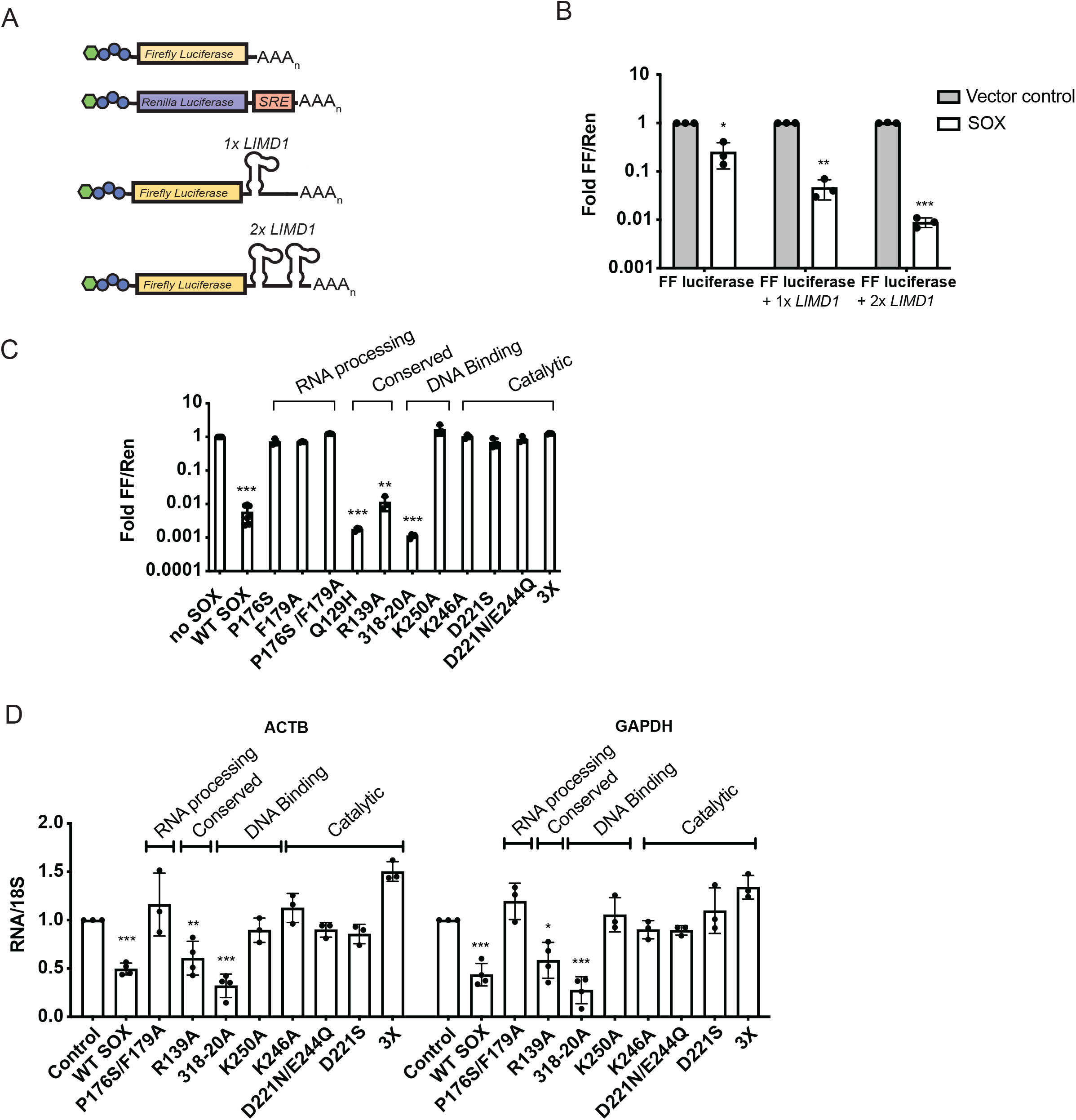
Catalytic residues, but not DNA binding activity, are required for the host shutoff function of SOX. (A) Schematic of the luciferase assay, in which firefly luciferase was engineered alone or with one or two SOX cleavage sites from the LIMD1 transcript. Renilla luciferase was engineered to contain the SOX resistant element (SRE) from the IL6 transcript rendering it deficient for SOX cleavage. (B) A dual luciferase assay was conducted to compare firefly luciferase signal normalized to Renilla luciferase signal in 293T cells transfected with the indicated constructs described in (A). Technical triplicate measurements were taken for each biological replicate. A minimum of three biological replicates were taken for each measurement. *P≤0.05, **P≤0.01, ***P≤0.001, one-sample t test versus hypothetical value of 1. (C) A dual luciferase assay was conducted using a firefly luciferase reporter containing a 2x LIMD1 SOX cleavage site as described in (B). Cells were transfected with empty vector or the indicated SOX mutants. A minimum of three replicate experiments are shown for each construct. **P≤0.01, ***P≤0.001, one-sample t test versus hypothetical value of 1. (D) The levels of cellular ACTB and GAPDH mRNA were measured by RT-qPCR in 293T cells transfected with the indicated SOX expression constructs or an empty vector control. All conditions were normalized to 18S rRNA values and then SOX-containing conditions are normalized to a vector control. Each dot is an independent biological replicate. *P≤ 0.05, **P≤0.01, ***P≤0.001, one-way ANOVA followed by Dunnett’s multiple comparisons test versus WT SOX.

We confirmed that these SOX mutants were expressed when transiently transfected into 293T cells (Supplementary Figure 5). Except for the 318-20 mutant, all mutants were expressed similarly to WT SOX. We also included SOX mutants P176S and F179A as controls, as they have previously been shown to be selectively required for SOX mRNA targeting (Supplementary Figure S5) (25, 42, 43). As expected, given that AEs generally use the same active site residues to process both DNA and RNA (35, 36), mutation of residues involved in catalysis on DNA (D221S, K246A, D221N/E224Q) greatly impaired SOX mRNA degradation (Figure 3C). The DNA processing defective mutants R139A and Q129H retained WT mRNA cleavage activity (Figures 3C), consistent with the previously description of Q129H as a separation of function mutant (11), However, the DNA binding mutant K250A was unable to target mRNA, suggesting it may contribute to both DNA and RNA binding, as indicated by the recent co-crystal structure of SOX bound to RNA (28) (Figures 3C). We confirmed the results obtained with the luciferase reporter by measuring the ability of the SOX mutants to degrade endogenous *ACTB* and *GAPDH* mRNAs in 293T cells, which yielded similar phenotypes (Figure 3D).

### SOX contributes to viral gene expression and virion production in iSLK cells

We next generated a KSHV BAC lacking SOX (SOX Stop) using the Red recombinase strategy, which allows for stable latent infection of KSHV mutants in the iSLK renal carcinoma cell line (28). These iSLK cells harbor a doxycycline-inducible version of the major KSHV lytic transactivator ORF50, allowing for efficient lytic reactivation of the BAC engineered virus (28). Because the ORF36 coding sequence overlaps the 5’ end of the SOX (ORF37) gene, we engineered two stop codons at amino acids 15 and 16 to maintain the full ORF36 coding sequence (Supplementary Figures S6A, S6B). We also generated a mutant rescue (MR) virus (SOX Stop MR) to verify that any observed phenotypes were due to the engineered change rather than secondary changes to the BAC sequence. We generated iSLK BAC16 cells harboring these constructs and confirmed the absence of SOX protein in the SOX Stop cell line by western blot and rescue of its expression in the SOX Stop MR cells (Figure 4A).

**Figure 4.**
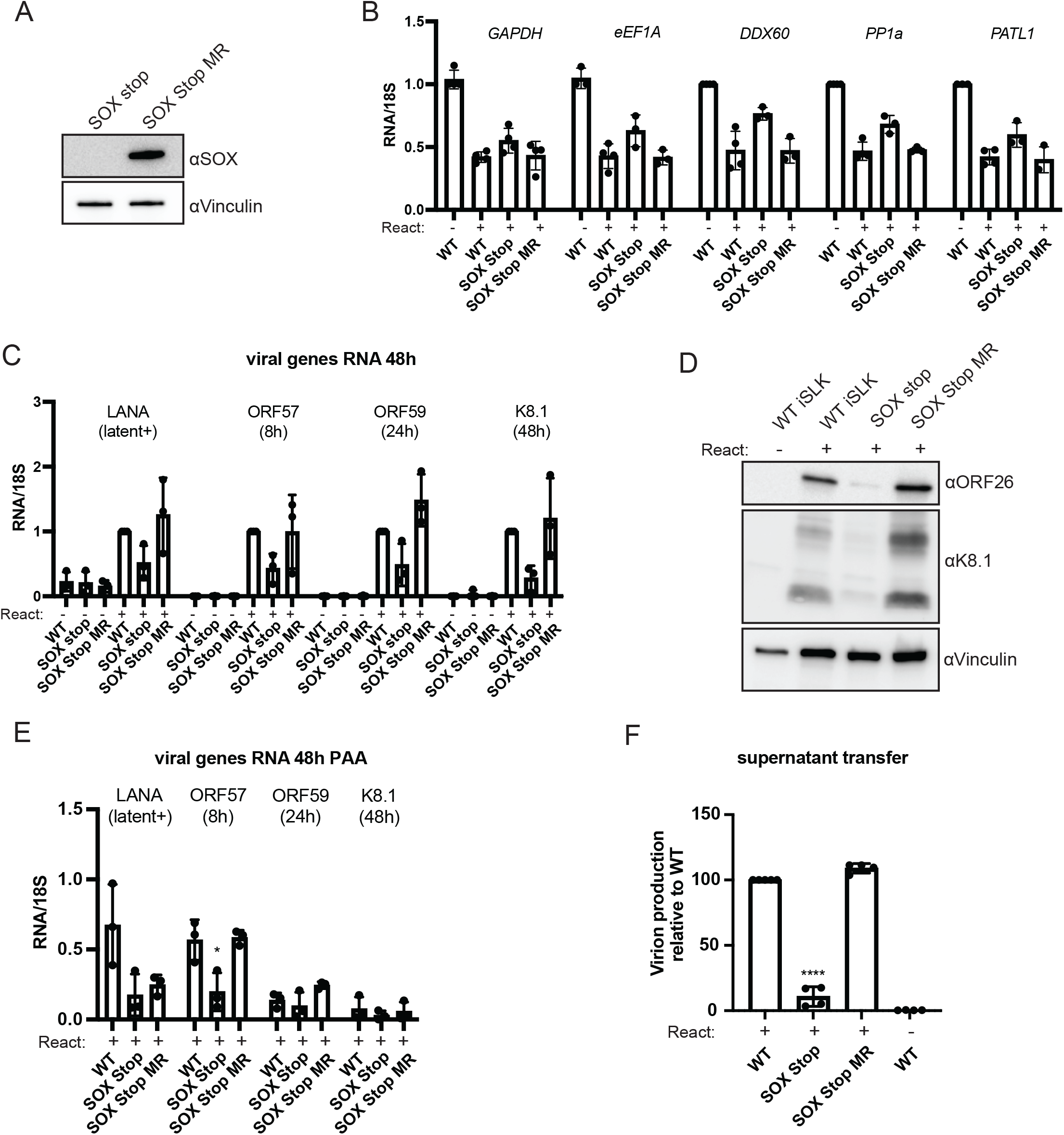
KSHV lacking SOX produces fewer infectious virions. (A)Western blot of whole cell lysates from iSLK cells containing a KSHV mutant that does not express SOX (SOX stop) or a mutant rescue version of KSHV where SOX expression is restored (SOX stop MR). Lysates were harvested 72 hours post lytic reactivation and blotted with an anti-SOX antibody. Vinculin serves as a loading control. (B-C) RT-qPCR was used to measure the RNA levels of several host (B) or viral (C) transcripts at 48 hours post-reactivation in iSLK cell lines containing the indicated WT or mutant KSHV. Transcript levels were normalized to 18S and to the WT iSLK unreactivated population for (B) for the WT iSLK reactivated population (C). A minimum of three biological replicates are shown. The peak timing of each of the viral genes’ expression is indicated below their name (52). (D) Western blot of lysates from the indicated iSLK cell lines that were either unreactivated or reactivated for 48h showing expression of late viral proteins ORF26 and K8.1. Vinculin is used as a loading control. (E) RT-qPCR was used as in (C) to measure the levels of the indicated viral transcripts, but in the presence of 500 mM of the viral DNA replication inhibitor phosphonoacetic acid (PAA). *P≤0.05, one-way ANOVA followed by Dunnett’s multiple comparisons test versus WT iSLK. (F) Infectious virion production was measured by flow cytometry on 1×106 293T cells infected with 1 mL of supernatant from 1×106 reactivated iSLKs. Bars represent the percentage of naïve 293T cells that became GFP positive after 24 hours. Data are from a minimum of four independent biological replicates. ****P≤0.001, one-sample t test versus hypothetical value of 1.

Given the dual roles of SOX as a DNase and RNase, we first sought to evaluate how the SOX Stop virus altered host mRNA levels. iSLK cells latently infected with WT KSHV, SOX Stop KSHV and SOX Stop MR were either untreated or lytically reactivated for 48 hours, whereupon mRNA levels were measured by RT-qPCR for 5 endogenous transcripts (*GAPDH, eEF1A, DDX60, PATL1, PP1a*) (Figure 4B). Reactivation of the WT or mutant rescue virus resulted in a 50% decrease of all five host transcripts, which was only partially rescued in the SOX Stop infection (Figure 4B). Thus, it is likely that other viral factors contribute in redundant ways to mRNA depletion in iSLK cells.

We next characterized the impact of SOX on other steps of the KSHV replication cycle. We evaluated viral gene expression by quantifying representative viral mRNAs from different kinetic classes by RT-qPCR at 48 hours post reactivation (Figure 4C). Relative to levels in WT KSHV or SOX Stop MR virus infections, the SOX Stop virus infection yielded lower mRNA expression for all viral transcripts tested. Measurements of protein levels of the late proteins ORF26 and K8.1 showed a similar trend where expression was lower in SOX Stop reactivated cells as compared to WT KSHV or SOX Stop MR (Figure 4D). We then tested whether the observed mRNA expression defects occurred downstream of viral DNA replication by treating the iSLK cell lines with phosphonoacetic acid (PAA), which pharmacologically inhibits the viral DNA polymerase (44). As expected, PAA treatment prevented expression of the DNA replication-dependent late viral transcript K8.1 in all the cell lines (Figure 4E). However, even in the presence of PAA, levels of ORF57 were significantly lower in the SOX Stop cells compared to the WT or SOX Stop MR cells (Figure 4E). Thus, the ability of SOX to potentiate expression of at least this early gene is independent of DNA replication. In contrast, there were less pronounced differences in the levels of LANA and ORF59 mRNA across the PAA-treated cell lines, suggesting that the reduction of these mRNAs in the SOX Stop cells may be primarily a downstream consequence of effects on DNA replication.

Finally, we assessed the cumulative effect of SOX function on the KSHV lifecycle by measuring production of infectious virions using a supernatant transfer assay. Supernatant from cells reactivated for 72 hours was incubated with naïve 293T cells and infectious virion production was determined by measuring the fraction of target 293T cells that became green from the GFP-marked virus. There was a 7-fold reduction in infection of target cells in SOX Stop relative to SOX Stop MR and WT (Figure 4F). Thus, SOX activity contributes to efficient KSHV amplification in iSLK cells, but it is not absolutely required for virion production in these cells.

### SOX DNA processing is dispensable for viral genome packaging but contributes to infectious virion production

A previous report has shown that SOX DNA processing activity does not affect intercellular levels of viral genomic DNA (11). However, we considered the possibility that its DNA processing activity influences whether the replicated DNA is competent for packaging into the viral capsid. We assessed viral genomic DNA packaging in lytically infected iSLK cells using a previously established packaging assay (33). During packaging, the viral genome is cleaved within the 20 to 40 copies of the terminal repeat (TR) sequence that separates unit-length genomes within concatemerized newly replicated DNA. The packaging motor cleaves stochastically, resulting in packaged viral genomes containing different numbers of TR sequences on either end. In this assay, non-TR DNA is removed by restriction digestion and the remaining ladder of TR concatemers formed during packaging are detected by Southern blotting.

We compared DNA packaging in KSHV containing WT SOX, the SOX stop mutant, the DNA processing mutant R139A, or the catalytic 3X mutant. Surprisingly, the TR cleavage pattern was similar for WT KSHV, SOX stop and the 3X mutant virus (Figure 5A), suggesting that SOX DNA processing activity is not required to prepare the viral genome for packaging. However, additional cleavage products were observed with the R139A and 3X mutant (Figure 5A, dashes), which may indicate their mutant activities could influence viral genome processing.

**Figure 5.**
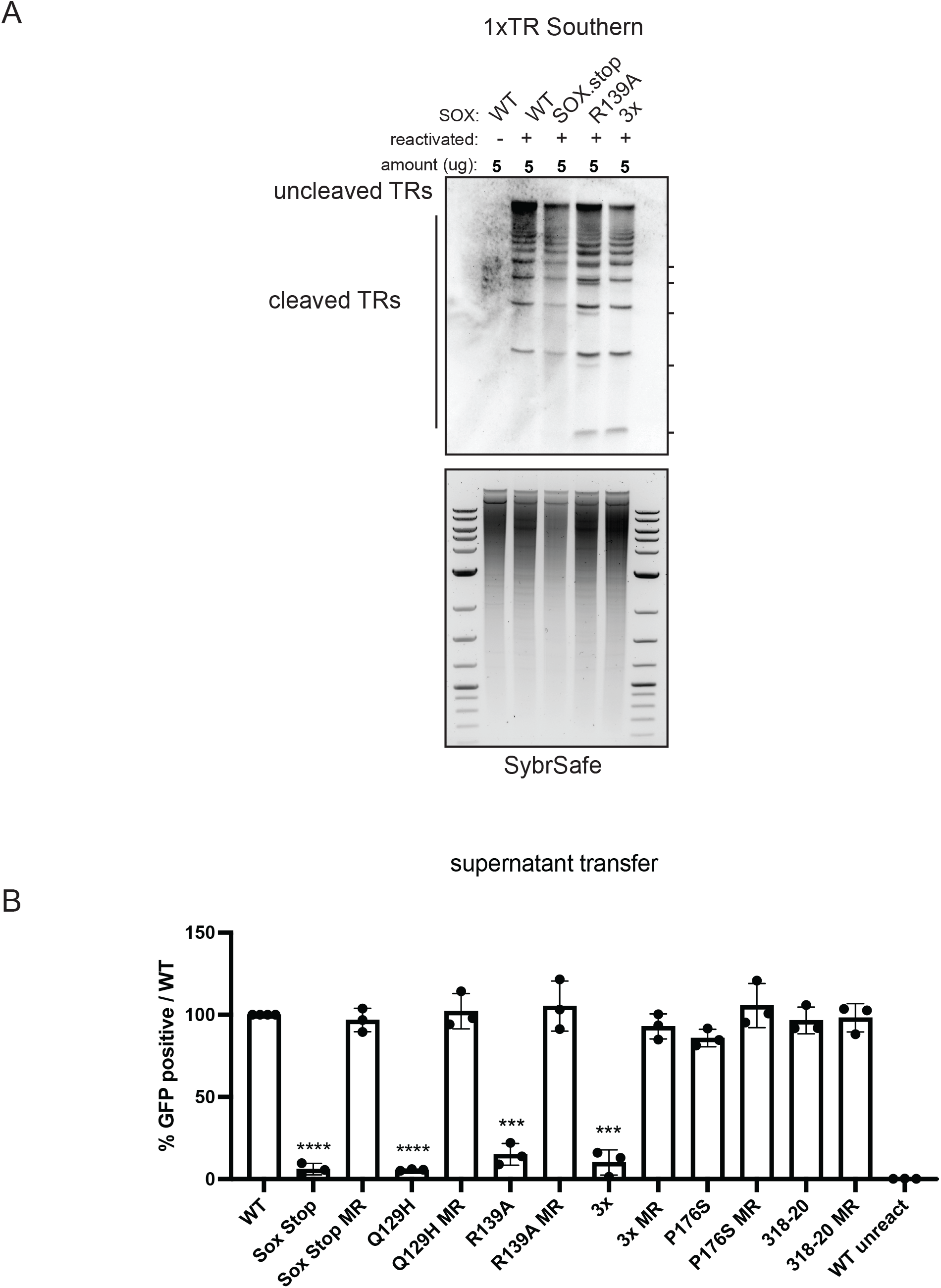
SOX residues involved in DNA processing are key for the KSHV lifecycle. (A) Southern blot of DNA isolated from iSLK cell lines using a probe for the terminal repeats. DNA was digested with PstI, which cuts within the genome but not within the terminal repeats (TR) and generates a ladder of (TR)-containing DNA when successful cleavage and packaging occurs. (B) Infectious virion production was measured by flow cytometry on 1×106 293T cells infected with 1 mL of supernatant from 1×106 reactivated iSLKs. Bars represent the percentage of naïve 293T cells that became GFP positive after 24 hours and are normalized to the WT condition. Data are from a minimum of three independent biological replicates. ***P≤0.001,****P≤0.001, one-sample t test versus hypothetical value of 1.

Finally, we leveraged our biochemical characterization of residues critical for the various SOX activities to determine which of its activities contribute to infectious virion production. In addition to the R139A and 3X catalytic mutant viruses described above, we used BAC16 engineering to generate KSHV mutants harboring the SOX Q129H mutant (selectively impaired for SOX activity on DNA) and the SOX P176S mutant (selectively impaired for processing mRNA). We also included the SOX 318-20 mutation, which has no apparent defect in any of the SOX activities we tested, as well as mutant rescue controls for each virus (Supplementary Figure S6C-D). In supernatant transfer assays, we observed that all three DNA processing mutants caused a significant reduction in KSHV virion production in comparison to their MR controls and to WT KSHV (Figure 5B). However, the RNA processing mutant P176S and the 318-20 mutant showed no defect in the production of infectious progeny virions (Figure 5B). Collectively, these data indicate that during iSLK infection, SOX-induced DNA processing plays a more important role than its mRNA cleavage activities (Figure 6).

**Figure 6.**
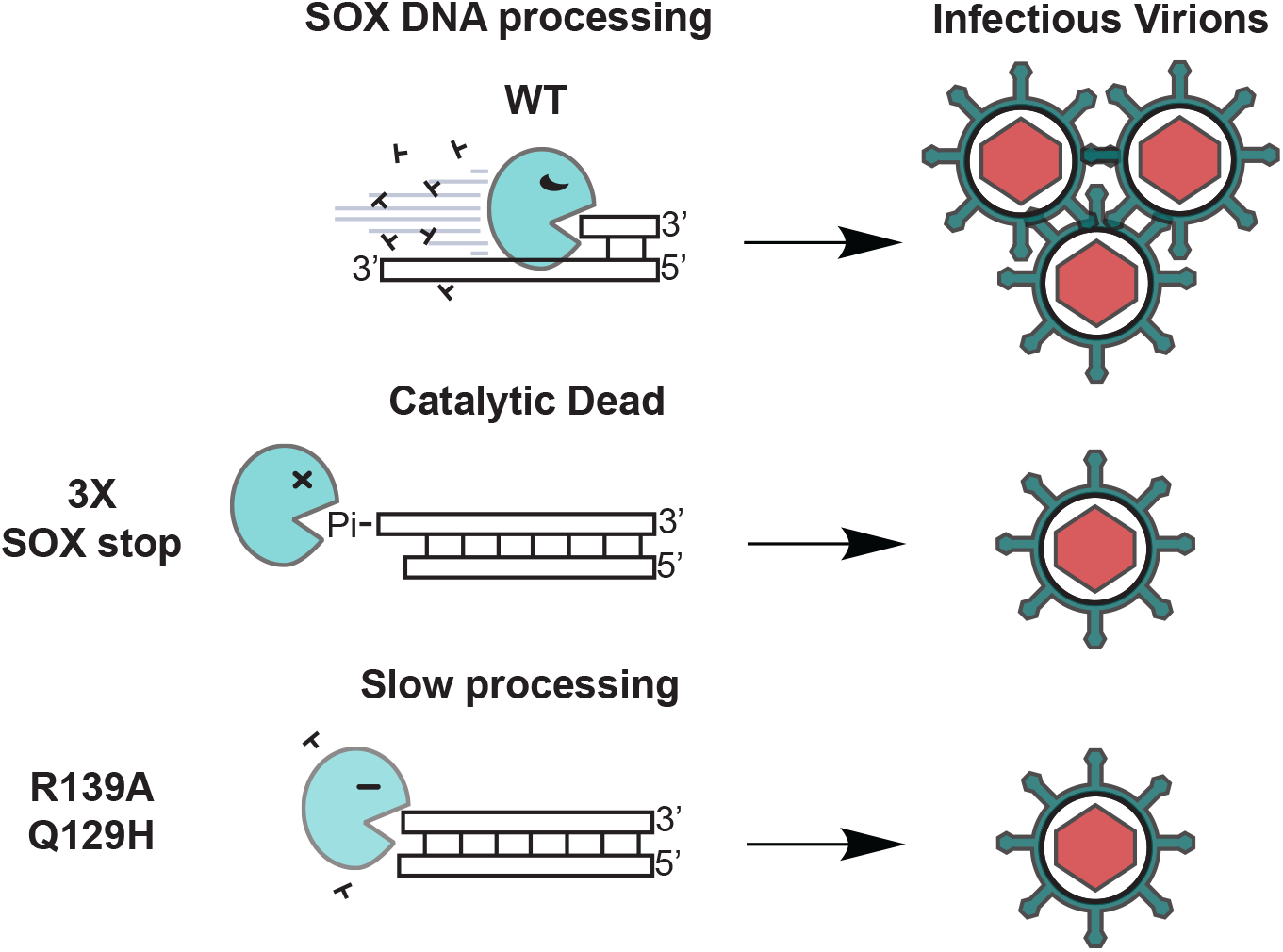
SOX end processing mutants produce fewer infectious virions. Cartoon representation of SOX end processing mutants resulting in the production of fewer infectious virions.

## Discussion

Multiple functional similarities have been found between herpesviruses and tailed bacteriophages, even though these families of dsDNA viruses infect different kingdoms of life (45). These similarities include replication of viral genomes in head-to-tail concatemers, the use of repeat sequences at genome termini for packaging into icosahedral capsids, and architecturally similar DNA packaging machinery (45, 46). Additional parallels in genome replication strategies are suggested by the fact that SOX and other herpesviral alkaline exonucleases are distantly related to the phage λ exonuclease (16). Our identification of residues positionally conserved between SOX and λ exonuclease that play central roles in SOX function underscores the evolutionarily ancient role of this enzyme in viral DNA replication and may further support the hypothesis that these viral lineages are derived from a common ancestor.

We observed that SOX binds DNA with nanomolar affinity in a manner independent of DNA sequence, in agreement with a co-crystal structure of SOX with DNA showing that it primarily interacts with the phosphate backbone (36). Interestingly, mutants R370 and 318-320 did not have any effect on SOX’s ability to interact with DNA. From the crystal structure, these residues participate in hydrogen bonding interactions with the phosphate backbone of DNA. It is possible that SOX could be bound in a non-productive or non-specific manner to the DNA substrate; such binding events have been observed with restriction endonucleases and with early crystal structures of lambda exonucleases (31, 47). In our DNA enzymatic assays we observe an inability of SOX to process non-phosphorylated blunt dsDNA which is the type of DNA substrate used in the co-crystal structure. It is likely that a complete understanding of how residues important for catalysis and end resection are engaging the 5’ end of DNA would be elucidated with a native DNA substrate. Furthermore, a comprehensive assignment of DNA binding residues would help to better understand how SOX peels off the 3’ end of DNA as it performs 5’-3’ exonucleolytic processing.

Our demonstration that specific residues in and surrounding the catalytic core of SOX contribute to completion of the viral lifecycle is well supported by data from multiple other herpesviruses (11, 42). Notably, several of the residues we characterized, including Q129 and R139 are highly conserved in all herpesvirus AEs as well as in the phage λ exonuclease. Neither Q129 nor R139 are part of the catalytic triad, but both are located within the active site and mutation of either is sufficient to significantly reduce SOX processing of DNA and impair virion production during infection. The observation that shorter DNA products accumulated upon mutation of Q129 and R139 suggests that these residues play a role in SOX processivity. Indeed, results from the phage λ exonuclease show a role for R139 in recognizing and processing the 5’ phosphate of DNA(31). Further structural studies trapping productive DNA and SOX binding intermediates would help to elucidate the role of other residues needed for processivity, including Q129.

We observed more limited SOX-induced changes to cellular mRNA abundance in iSLK cells than has been reported for KSHV infection of microvascular endothelial cells or infection of multiple cell types with the related g-herpesvirus MHV68 (20, 22). This could be due to a stronger contribution of other viral factors to host shutoff in iSLK cells, or possibly to reduced expression of a host cofactor(s) involved in SOX RNase activity. This parallels cell-type specific findings with the HSV-1 host shutoff factor vhs in which viral genes are translated more poorly during infection with a vhs null virus in several (e.g., HeLa, HepG-2) but not all (e.g., Vero) cell lines (48). Indeed, viral activities central to genome replication should have prominent phenotypes in all cell types whereas the contributions of virus-host interactions that influence immune evasion or optimize gene expression could vary by cell type. The mRNA targeting function of SOX, which has established importance *in vivo*, presumably falls in this latter category (22).

The precise role of SOX and other AEs in viral DNA replication remains an open question, but several hypotheses have been put forth based on observations from HSV-1 and phage λ. Branched structures accumulate in herpesvirus infection in the absence of AEs, suggesting they play a role in resolving these structures (9, 10, 49). The altered KSHV DNA banding pattern in the packaging assay with the R139A and catalytic 3X SOX mutants could be indicative of alternative forms of the viral DNA that arise from inefficient branched structure resolution. Indeed, herpesviral AE mutants have been shown to impair accumulation of DNA-containing capsids, including upon infection with KSHV containing the Q129H SOX DNA processing mutant (7–9, 11, 49). While our results show that SOX activity is not essential for DNA cleavage during packaging, we cannot exclude the possibility that it influences packaging, as the cleavage assay does not quantitatively measure packaging efficiency. The packaged DNA in virions lacking functional SOX could also be less ‘intact’ or lower quality than in WT KSHV, which would result in the observed reduction in the number of infectious particles produced.

The HSV-1 AE has also been hypothesized to promote viral genome replication by inducing a single strand annealing form of recombination (50). We see that KSHV SOX has the greatest activity on ss and dsDNA substrates with a 5’ phosphate, consistent with a role in processing 5’ overhangs that could facilitate recombination. Perhaps analogous to the λ Red recombination system, this could occur if SOX continued to partially or fully degrade the overhanging strand of the DNA duplex to create the ssDNA template needed for strand invasion or single-stranded 3’ end annealing. Annealing of the 3’ ssDNA end to its homologous end is catalyzed by Redβ, a ssDNA binding protein and binding partner of λ exonuclease (51). In HSV-1, the AE (UL12) interacts with the viral single stranded DNA binding protein ICP8 to promote strand exchange (15, 50), like a Redα/Redβ system. A key remaining question for herpesviruses is how an AE would both promote strand invasion and resolve complex DNA structures.

Finally, the SOX-linked defects in viral gene expression could reduce infectious virion production by negatively impacting virion egress. Previous work in EBV has shown that a virus lacking the SOX homolog BGLF5 has defects in nuclear egress, thereby resulting in an accumulation of empty C capsids (9). The mutants characterized herein should prove valuable for future experiments to delineate the contributions of each of the biochemical activities of SOX on different facets of the viral lifecycle.

## Acknowledgments

We thank all members of the Glaunsinger and Coscoy labs for their helpful suggestions and discussions. This work was funded by NIH grant CA136367 to B.G., who is also an investigator of the Howard Hughes Medical Institute. E.H. is supported by a NSF GFRP fellowship. A.L.D. was supported by the Rhee Family Fellowship of the Damon Runyon Cancer Research Foundation (DRG-2349-18).

