## Supplemental figures and tables for "DNA processing by the Kaposi’s sarcoma-associated herpesvirus alkaline exonuclease SOX contributes to viral gene expression and infectious virion production"

### Supplementary Figure 1

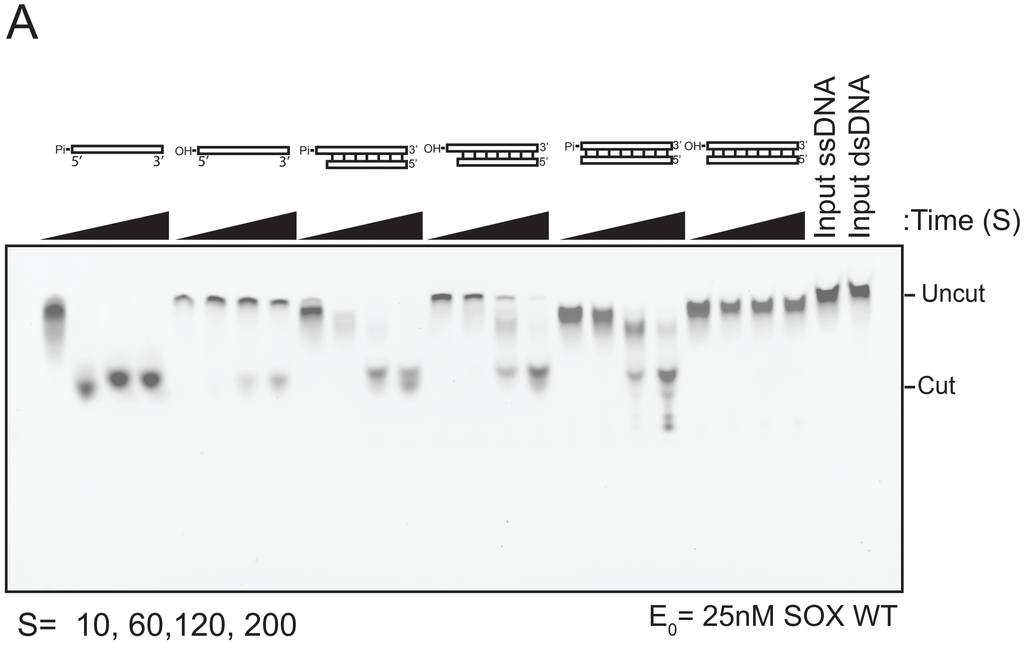


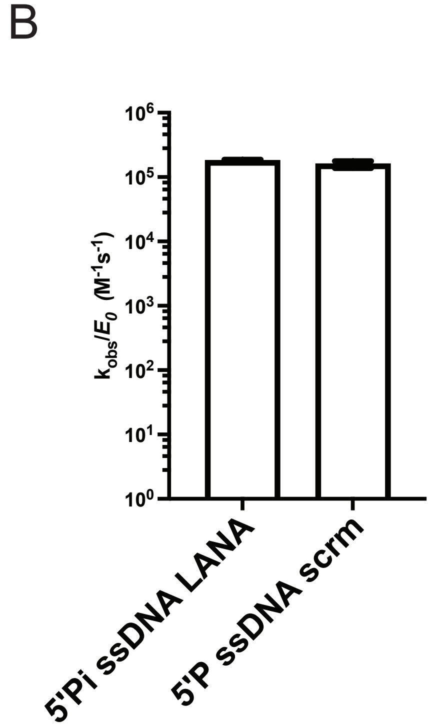


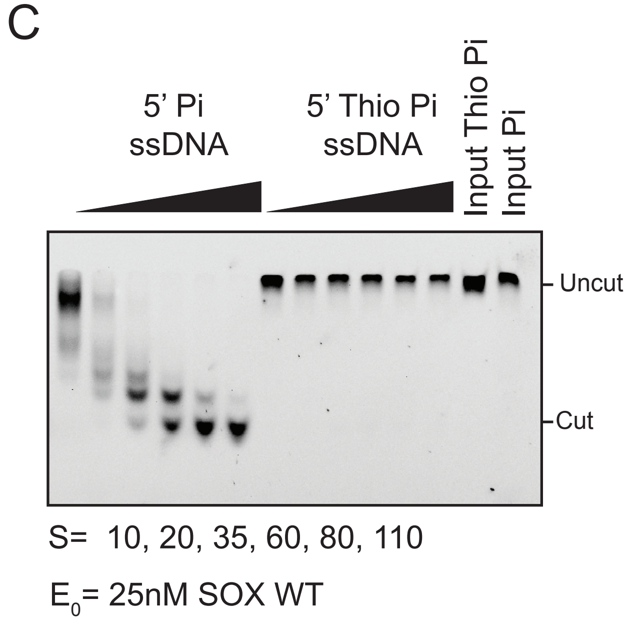


**Supplementary Figure 1**
(A) Urea-PAGE gel showing DNA processing by WT SOX of the dsDNA and ssDNA substrates used in Figure 1B. DNA substrates contain either a 5' phosphate (Pi) or hydroxyl (OH) group. All substrates contain a 3' fluorophore, which is located on the antisense strand for dsDNA substrates. Reactions were carried out at concentration of 25 nM SOX and time was recorded in seconds (s). (B) The catalytic efficiency of SOX processing of a 25 bp ssDNA LANA and a 25 bp ssDNA scrambled sequence. Reactions were run at a constant concentration of 25 nM. (C) Urea-PAGE gel showing WT SOX (25 nM) processing of a 5' phosphate containing DNA probe under substrate limiting conditions, and inhibition of processing by the 5' thio-phosphate DNA probe. Time points are shown below the gel. Markers indicate cut and uncut DNA products.

### Supplementary Figure 2

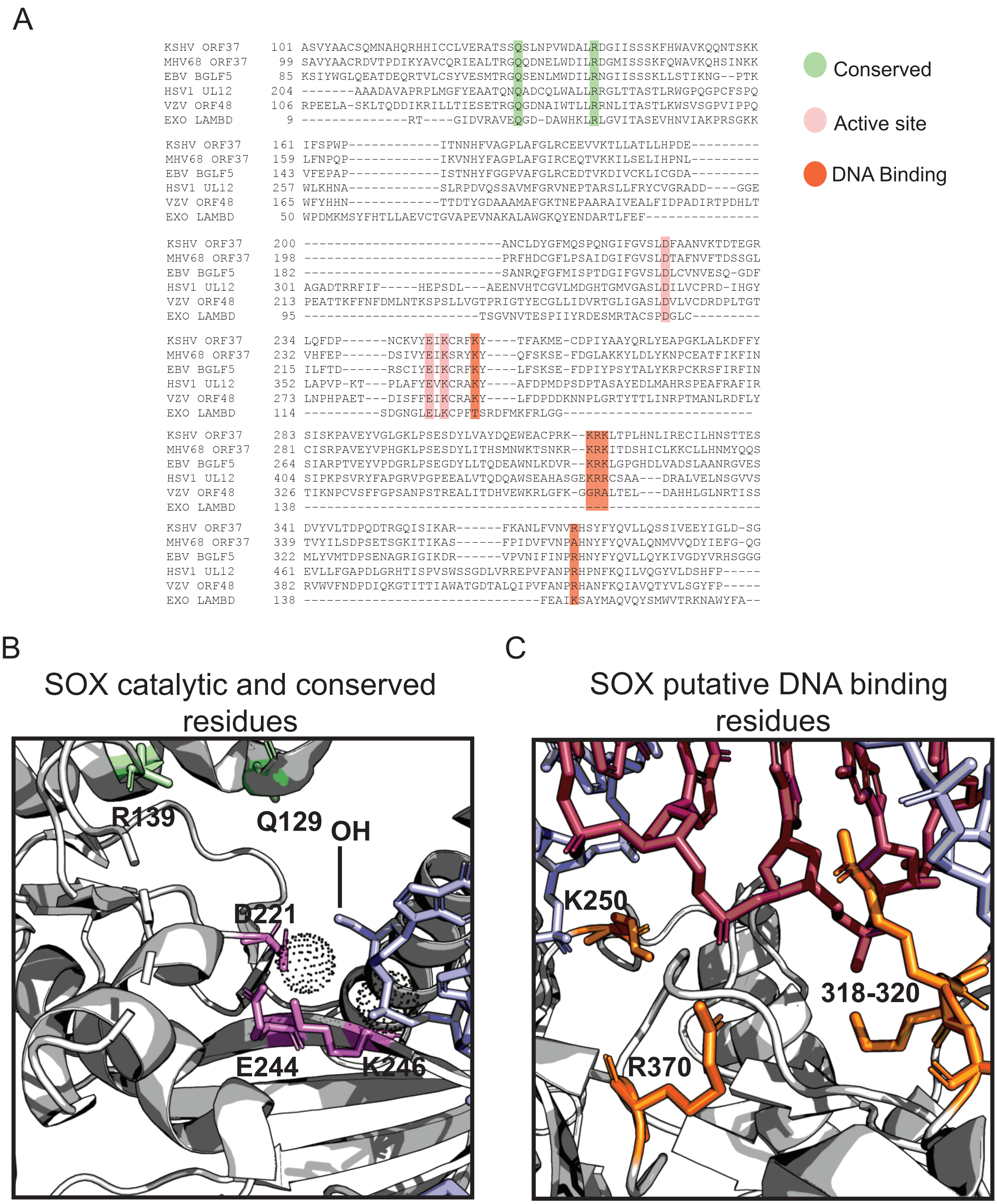


**Supplementary Figure 2**
(A) Sequence alignment using Expasy uiprot alignment tool of herpesviral exonucleases spanning alpha- and gamma-herpesvirus subfamilies as well as λexonuclease. Conserved residues are highlighted in green. Residues highlighted in orange contact DNA in PDB:3POV structure. Residues highlighted in pink indicate residues predicted to be part of the active site. (B) Structure of the KSHV SOX active site. Catalytic residues are highlighted pink, conserved residues highlighted in green and spotted spheres are magnesium site 1 and site 2. The 5' hydroxyl DNA is colored light blue and positioned in the active site (PDB:3POV). (C) Putative DNA binding residues are highlighted in orange. Double stranded DNA (light blue 5' and cyan 3') is modeled into SOX (34).

### Supplementary Figure 3

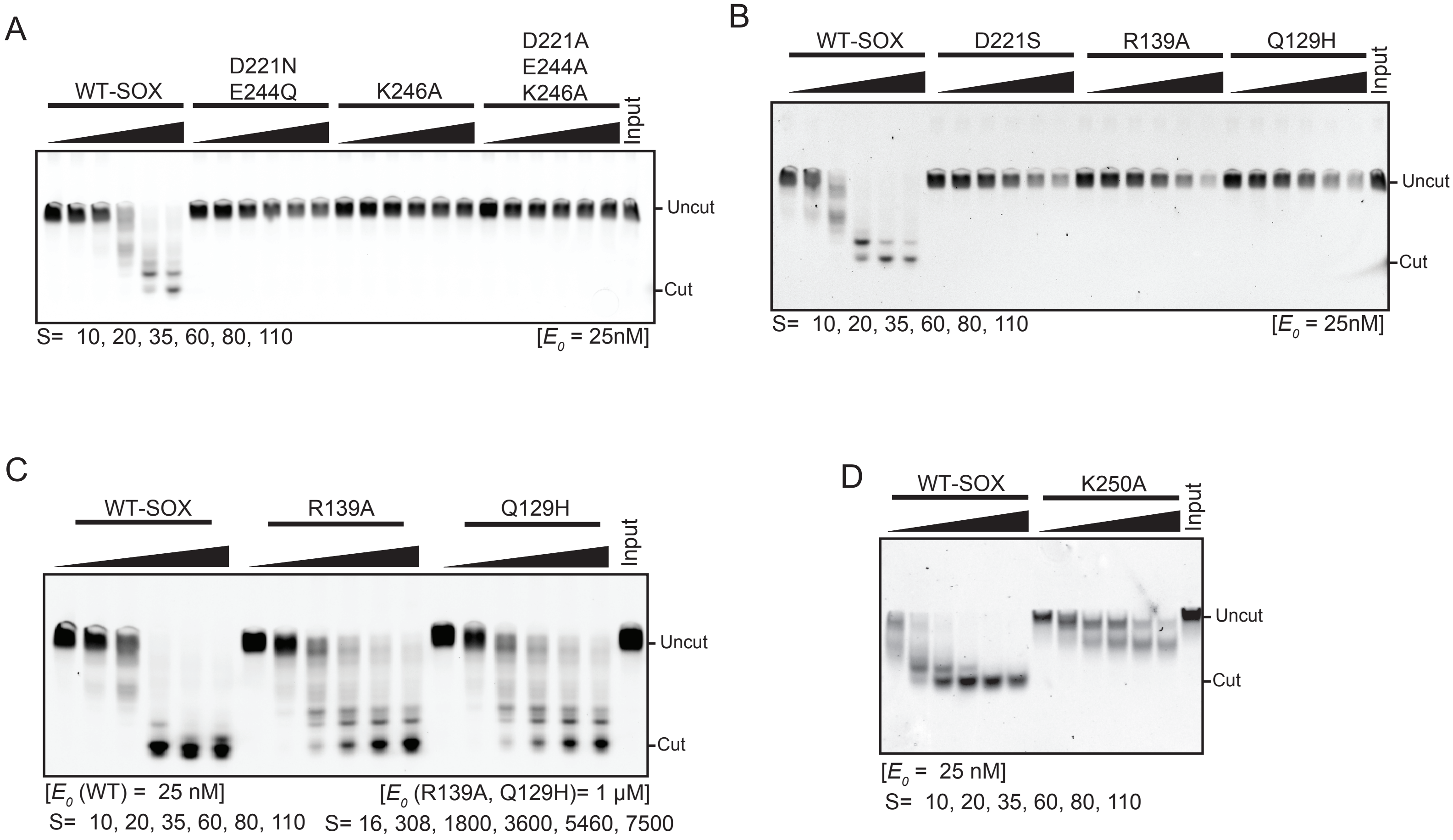


**Supplementary Figure 3**

(A) Urea-PAGE gel showing DNA processing by WT SOX and the D221N/E224Q, K246A and D221A/E244A/K246A (3X) SOX mutants. The reactions were run at 25 nM of enzyme and hash marks denote uncut and cut DNA substrate. Times listed below the gel indicate points at which the reactions were stopped. (B) Urea-PAGE gel showing DNA processing by WT, D221S, R139A and Q129H SOX. The reactions were run at 25 nM of enzyme and hash marks denote uncut and cut DNA substrates. Times listed below the below the gel indicate points at which the reactions were stopped. (C) Urea-PAGE gel showing processing defects of SOX mutants in conserved residues R139A and Q129H. The reactions were run at 1 uM of R139A and Q129H, and 25 nM of WT SOX enzyme. Hash marks denote uncut and cut DNA substrates. Times listed below gel indicate points at which the reactions were stopped. (D) Urea-PAGE gel showing DNA processing by K250A (25 nM) and WT SOX (25 nM). Hashmarks denote uncut and cut DNA substrates. Times listed below gel indicate points at which the reactions were stopped.

### Supplementary Figure 4

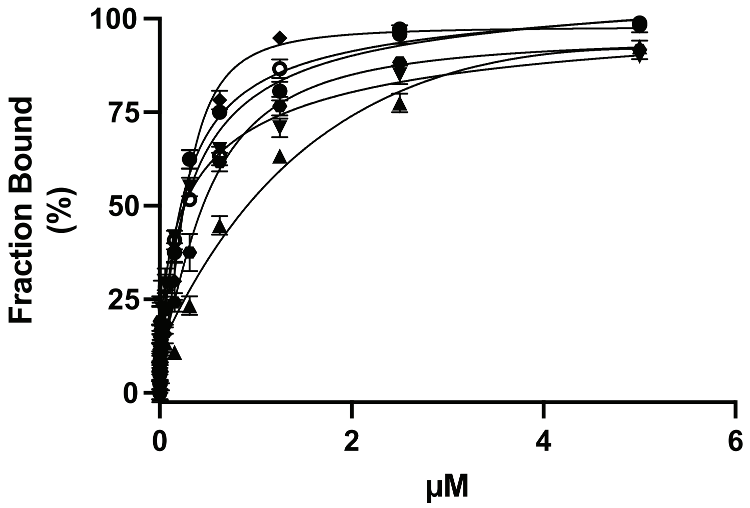


| SOX | Affinity (nM) |
| --- | --- |
| WT | 143.45 ± 33.45 |
| Q129H | 105.52 ± 40.15 |
| K250A | 649.12 ± 39.25 |
| R139A | 189.32 ± 49.25 |
| R370A | 120.40 ± 45.5 |
| 318-320A | 120.39 ± 39.5 |

**Supplementary Figure 4**
(A) Equilibrium binding measurements of WT SOX (hollow circle), Q129H (solid circle) K250A (triangle), R139A (diamond), R370A (square), 318-320A (upside down triangle). A constant concentration (1nM) of a fluorescently labeled LANA probe was used as described in Figure 1A. Data represent 3 biological replicates. The table shows calculated Kd values for SOX proteins using a signal binding model.

### Supplementary Figure 5

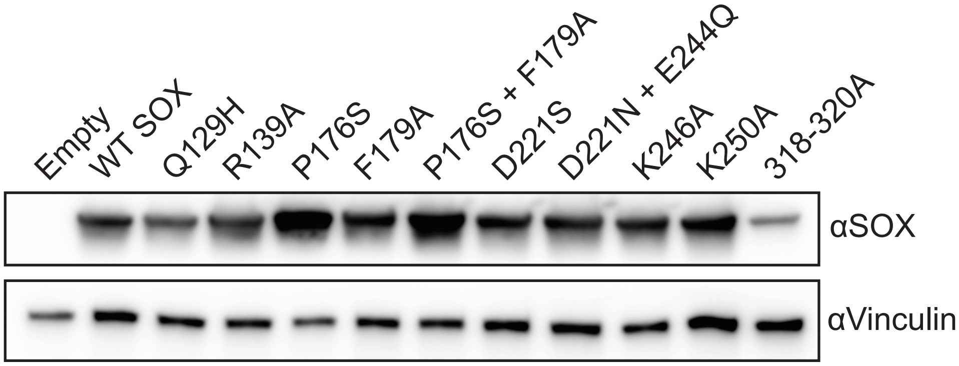


**Supplementary Figure 5**
(A) Western blot of 293T cell lysates transiently transfected with the indicated SOX mutant expression constructs and probed with an anti-SOX antibody. Vinculin serves as a loading control.

### Supplementary Figure 6

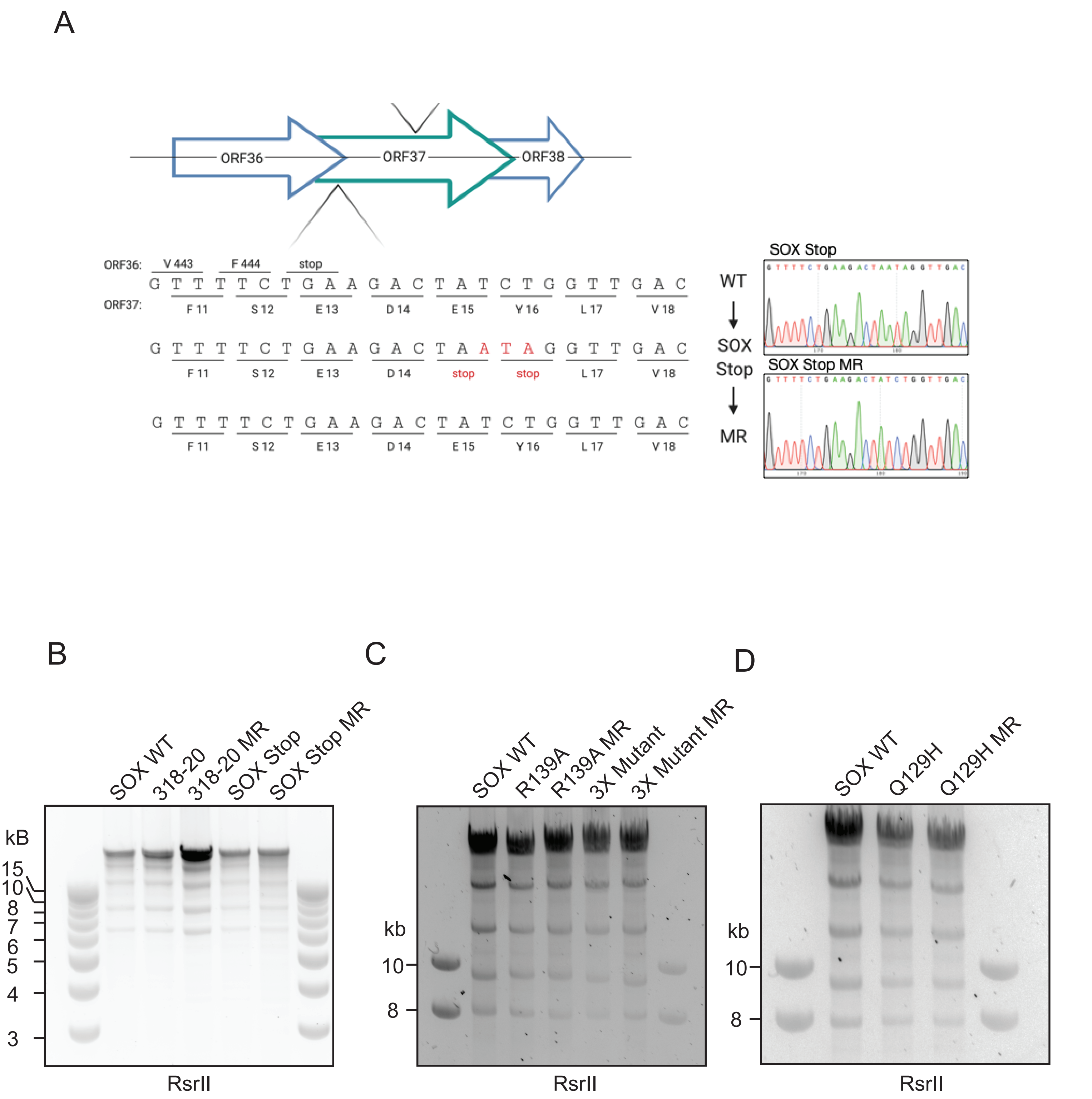


**Supplementary Figure 6**
(A) Diagram depicting the genomic locus of the engineered SOX Stop mutant, shown in red. Sanger sequencing for SOX Stop and the MR cell line at the site of mutation is shown adjacent to the diagram. (B-D) RsrII digestions of the engineered BACs in comparison to the WT Bac16 to demonstrate no large-scale rearrangements in BAC DNA.

### Supplementary table 1

| mutant | function | dsDNA catalytic efficiency M-1/s-1 | reference |
|---|---|---|---|
| WT SOX | | 1.78x10^6 +/- 4.3x10^3 | this paper |
| P176S* | RNA processing | N/A | (25) |
| F179A* | RNA processing | N/A | (25) |
| K250A | DNA binding | 1.64X10^5 +/- 6.5x10^3 | this paper |
| 318-20A | DNA binding | 1.47X10x^6 +/- 6.1X10^4 | this paper |
| R370A | DNA binding | 1.24x10^6+/- 9.1x10^4 | this paper |
| D221S | Catalytic residue | N/D | this paper |
| K246A | Catalytic residue | N/D | this paper |
| D221N/E244Q | Catalytic residue | N/D | this paper |
| D221N+E244A+K246A 3X mutant | Catalytic residues | N/D | this paper |
| Q129H | conserved | 1.0x10^3 +/- 987 | (11) |
| R139A | conserved | 1.1x10^3 +/- 785 | this paper |

* Previously characterized

**Supplementary table 1**
Table of SOX mutants used in this study are listed by functional group, the indicated function of the mutated residue is noted and their catalytic efficiency on ssDNA is annotated, if relevant, with indication of the publication that characterizes their activity. ND is non detected and N/A is not applicable as these residues are involved in RNA processing.

### Supplementary table 2

| | |
|---|---|
| 18S fwd | GTAACCCGTTGAACCCCATT |
| 18S rev | CCATCCAATCGGTAGTAGCG |
| ACTB cds fwd | CATGTACGTTGCTATCCAGGC |
| ACTB cds rev | CTCCTTAATGTCACGCACGAT |
| GAPDH cds fwd | CTGGGCTACACTGAGCACC |
| GAPDH cds rev | AAGTGGTCGTTGAGGGCAATG |
| PP1A cds fwd | TCCTGGCATCTTGTCCATG |
| PP1A cds rev | CCATCCAACCACTCAGTCTTG |
| CTGF promoter fwd | CGAGGAATGTCCCTGTTTGT |
| CTGF promoter rev | ACTGGCTGTCTCCTCTCAGC |
| DDX6 cds fwd | CAGGAACATCGAAATCGTG |
| DDX6 cds rev | TCCAATACGATGGAGATAGG |
| PATI1 cds fwd | TCCTGCTCCCTATGGTGAGAG |
| PATI1 cds rev | CATGGCAGCAAGTGGACTACC |
| KSHV ORF59 promoter fwd | AATCCACAGGCATGATTGC |
| KSHV ORF59 promoter rev | CACACTTCCACCTCCCCTAA |
| KSHV ORF57 cds fwd | GGTGTGTCTGACGCCGTAAAG |
| KSHV ORF57 cds rev | CCTGTCCGTAAACACCTCCG |
| KSHV ORF59 cds fwd | GAAGGCAGTGGAGACGTTAGC |
| KSHV ORF59 cds rev | ACGGTGGAGGTGAGGTTGTC |
| KSHV K8.1 cds fwd | CCGTCGGTGTGTAGGGATAAAG |
| KSHV K8.1 cds rev | GTCGTTGTAGTGGTGGCAGAAA |
| KSHV ORF31 cds fwd | CAACATTCAAGAGGGCCAGT |
| KSHV ORF31 cds rev | ATGTCTGCAAATACCGCACA |
| KSHV ORF66 cds fwd | CCCATGCCACTTCTTCCTTA |
| KSHV ORF66 cds rev | CCACGTATGGGCAAAATACC |
| KSHV LANA cds fwd | TGGAGATGGGAGATGTAGGC |
| KSHV LANA cds rev | TTCGCCAACCGTAGTGTGTA |

**Supplementary table 2**
Table of primers used in this publication.
